# Identification of a novel *Candida metapsilosis* isolate suggests ongoing hybridization

**DOI:** 10.1101/2021.07.15.452539

**Authors:** Caoimhe E. O’Brien, Bing Zhai, Mihaela Ola, Eoin Ó Cinnéide, Ísla O’Connor, Thierry Rolling, Edwin Miranda, N. Esther Babady, Tobias M. Hohl, Geraldine Butler

## Abstract

*Candida metapsilosis* is a member of the *C. parapsilosis* species complex, a group of opportunistic human pathogens. Of all the members of this complex, *C. metapsilosis* is the least virulent, and accounts for a small proportion of invasive *Candida* infections. Previous studies established that all *C. metapsilosis* isolates are hybrids, originating from a single hybridization event between two lineages, parent A and parent B. Here, we use MinION and Illumina sequencing to characterize a *C. metapsilosis* isolate that originated from a separate hybridization. One of the parents of the new isolate is very closely related to parent A. However, the other parent (parent C) is not the same as parent B. Unlike *C. metapsilosis* AB isolates, the *C. metapsilosis* AC isolate has not undergone introgression at the Mating Type-like Locus. In addition, the A and C haplotypes are not fully collinear. The *C. metapsilosis* AC isolate has undergone Loss of Heterozygosity (LOH) with a preference for haplotype A, indicating that this isolate is in the early stages of genome stabilization.

## INTRODUCTION

*Candida metapsilosis* is a rare opportunistic pathogen of humans (Gomez-Lopez *et al*. 2008; Lockhart *et al*. 2008a; Silva *et al*. 2009). It is a member of the *Candida parapsilosis* species complex, a small clade of related organisms that includes *C. metapsilosis, Candida orthopsilosis* and *C. parapsilosis sensu stricto*, all of which cause infection in humans (Tavanti *et al*. 2005). Within this group, *C. parapsilosis* is the most common cause of candidiasis, whereas *C. metapsilosis* is the least common, with an incidence ranging from 0.6 -6.9% of cases of invasive candidiasis (Gomez-Lopez *et al*. 2008; Lockhart *et al*. 2008a; Silva *et al*. 2009; Cantón *et al*. 2011; Bonfietti *et al*. 2012; Bertini *et al*. 2013). However, isolates from the *C. parapsilosis* species complex are commonly misidentified, which may have led to an underestimation of the frequency of *C. orthopsilosis* and *C. metapsilosis* (Tavanti *et al*. 2007; Lockhart *et al*. 2008b; Bonfietti *et al*. 2012). In recent years, it has been suggested that the incidence of *C. orthopsilosis* and *C. metapsilosis* infection is increasing, although this may be due to increased awareness of the species differentiation (Lockhart *et al*. 2008a).

Few *C. metapsilosis* isolates secrete virulence-associated factors such as lipases or aspartic proteinases in comparison to *C. parapsilosis* (Németh *et al*. 2013). Whereas *C. parapsilosis sensu stricto* is commonly associated with infections in neonates, *C. metapsilosis* is rarely associated with neonatal infection and appears to affect adults predominantly (Cantón *et al*. 2011). There is no evidence of widespread resistance to any antifungal drugs in *C. metapsilosis* and most isolates tested thus far are susceptible to antifungals (Gomez-Lopez *et al*. 2008). There is some suggestion that *C. metapsilosis* may be a human commensal, and it has been isolated from the oral cavity of healthy individuals (Ghannoum *et al*. 2010).

Although all members of the *C. parapsilosis* species complex have diploid genomes, *C. parapsilosis* isolates are highly homozygous with, on average, 0.06 heterozygous SNPs per kb (Butler *et al*. 2009; Pryszcz *et al*. 2013), whereas the majority of isolates of *C. orthopsilosis* and *C. metapsilosis* are extremely heterozygous (Pryszcz *et al*. 2014; Schröder *et al*. 2016). Most *C. orthopsilosis* isolates have heterozygosity levels ranging from 8 -31 SNPs per kilobase, and originated from multiple hybridization (mating) events between related parents (Pryszcz *et al*. 2014; Schröder *et al*. 2016). Previous analysis of 11 *C. metapsilosis* isolates showed that they were highly heterozygous, ranging from 22 -26 SNPs per kilobase (Pryszcz *et al*. 2015). The authors proposed that these *C. metapsilosis* isolates arose from hybridization between two parental lineages that differed by approximately 4.5% divergence at the genome level.

Ten of the 11 *C. metapsilosis* isolates previously analyzed by (Pryszcz *et al*. 2015) are heterozygous at the Mating Type-like Locus, with both *MTL***a** and *MTLalpha* idiomorphs present. The *MTLalpha* locus is intact, and is identical in the arrangement and orientation of its genes to the *MTLalpha* locus in *C. albicans, C. tropicalis* and *C. orthopsilosis* (Pryszcz *et al*. 2015). However, introgression has occurred at the *MTL***a** locus, where the *PAP***a**, *OBP***a** and *PIK***a** genes present in most *Candida* species have been overwritten with *MTLalpha*2, *OBPalpha*, and a portion of *PIKalpha*. Because the introgression is present in almost all sequenced *C. metapsilosis* isolates, it is likely that hybridization occurred once, followed by introgression, and all extant isolates descended from this. The 11th *C. metapsilosis* isolate is missing all of *MTL***a**, which Pryszcz et al. (Pryszcz *et al*. 2015) proposed resulted from an additional LOH event that has overwritten the remainder of the cassette.

Many fungal species that infect humans are hybrids, including *C. orthopsilosis* (Pryszcz et al. 2014; Schröder et al. 2016), *Candida inconspicua* (Mixão et al. 2019) and *C. tropicalis* (O’Brien *et al*. 2021). In some fungal pathogens, such as *Cryptococcus neoformans* (Li et al. 2012), and the plant pathogen *Verticillium longisporum* (Inderbitzin *et al*. 2011), hybridization is associated with increased virulence, increased antifungal resistance or an expanded host range (reviewed in (Mixão and Gabaldón 2018)). In *C. orthopsilosis* (Schröder *et al*. 2016) and *Cryptococcus neoformans* (Xu *et a*l. 2002; Li *et al*. 2012), multiple hybridization events have occurred and may be ongoing. Here, we describe the discovery of a novel hybrid of *C. metapsilosis* isolated from human feces. This isolate originated from a hybridization event between one parent that is similar to one of the parents of the previously sequenced isolates, and a second parent that is approximately 4.5% different. We therefore propose that hybridization is also ongoing in the *C. metapsilosis* lineage.

## MATERIALS AND METHODS

### DNA extraction and Illumina sequencing

The isolates used in this study are shown in Table S1. Strains from Memorial Sloan Kettering Cancer Center were cultured on Sabouraud (SAB) agar for 48 h at 37°C, then grown in overnight culture in 2 -3 ml of Yeast Extract-Peptone-Dextrose (YPD) broth at 240 rpm. Genomic DNA was extracted and DNA libraries were sequenced on an Illumina HiSeq platform generating 100 bp paired-end reads, as described in Zhai et al. (Zhai *et al*. 2020). Some isolates were previously described in Zhai et al. (Zhai *et al*. 2020). Illumina data from *C. metapsilosis* strain ATCC 96143 were downloaded from the NCBI Sequence Read Archive (SRA) under the BioProject ID PRJNA432377 (Oh *et al*. 2019). Illumina reads from (Pryszcz *et al*. 2015) were downloaded from the SRA under BioProject ID PRJEB1698. The quality of all Illumina data was checked using FastQC (https://www.bioinformatics.babraham.ac.uk/projects/fastqc/). Reads were trimmed with Skewer version 0.2.2, using parameters “-m pe” (paired end mode) “-l 50” (minimum read length allowed after trimming is 50 bases) “-q 15” (trim 3’ end until quality of 15 is reached) “-Q 15” (lowest mean quality allowed before trimming) (Jiang *et al*. 2014). Data were assembled using SPAdes version 3.13.1 with the --careful parameter (Bankevich *et al*. 2012).

### MinION sequencing

*C. metapsilosis* MSK414 was cultured on YPD agar for 48 h at 30°C, then grown overnight in 50 ml YPD broth at 30°C, 200 rpm. DNA was extracted using the QIAGEN Genomic-Tip (100/G) kit as per kit instructions. DNA quality was checked with NanoDrop and quantified using the Qubit fluorometer. Libraries were prepared for minION sequencing and barcoded with the Rapid Barcoding Kit (SQK-RBK004) from Oxford Nanopore Technologies (ONT). Prepared libraries from three species were pooled and loaded onto an ONT minION flow cell (FLO-MIN106) for sequencing for 50 h. Basecalling for the minION data was performed using the ONT Guppy software, version 3.2.4+d9ed22f with the following parameters; “--input_path fast5 --save_path fastq --flowcell FLO-MIN106 --kit SQK-RBK004 --verbose_logs --cpu_threads_per_caller 5 --num_callers 7”. Basecalled data were demultiplexed using qcat version 1.1.0 with the following parameters; “--fastq fastq/all_multiplexed_reads.fastq --barcode_dir demultiplex_qcat --detect-middle --min-read-length 1 --trim --kit RBK004 --epi2me”. After demultiplexing, 2.79 Gb of data were assigned to *C. metapsilosis* MSK414. Read quality was checked with NanoPlot version 1.23.1 (De Coster *et al*. 2018). Potential contaminant reads (i.e. any reads mapping to the other species sequenced in multiplex on the flow cell) were identified by command-line BLASTN from the BLAST+ package version 2.2.31 (Camacho *et al*. 2009) and reads with Q < 7 and length < 1 kilobase (kb) were removed with NanoFilt version 2.3.0 (De Coster *et al*. 2018). Post-filtering, 2.76 Gb of data assigned to *C. metapsilosis* MSK414 were available for analysis.

Filtered MinION reads were assembled using Canu version 1.8 using recommended parameters for haplotype separation; -p canu_run2 -d canu_run2 genomeSize=13511817 corOutCoverage=200 “batOptions=-dg 3 -db3 -dr 1 -ca 500 -cp 50” -nanopore-raw all_q7l1k.fastq (Koren *et al*. 2017). Illumina reads were used to polish the assembly with Pilon version 1.23 (Walker *et al*. 2014). Assembly statistics were checked using QUAST version 4.6.1 (Gurevich *et al*. 2013). The final assembly consisted of 45 contigs, with 18 chromosomal-sized contigs, an N50 of 1.78 Mb, and an L50 of 6 (Table S2). Zeros were removed from the beginning of contig names for clarity. Circos version 0.69 (Krzywinski et al. 2009) and Circoletto (Darzentas 2010) were used to visualize alignments.

Each contig in the *C. metapsilosis* MSK414 assembly was assigned to either the A or C parent based on its percentage identity to the best hit in the assembly of *C. metapsilosis* ATCC 96143 (Oh *et al*. 2019). Global percentage identity was measured using MUMmer dnadiff version 1.3 with default options (Kurtz et al. 2004). Pairs of contigs mapping to one contig in the *C. metapsilosis* ATCC 96143 reference assembly were assigned as alternative haplotypes of the same chromosome.

### Variant calling and filtering

Variants were called from the Illumina data. Trimmed reads were aligned to the chimeric reference assembly produced by (Pryszcz *et al*. 2015). Reads were aligned using bwa mem version 0.7.12-r1039 with default parameters (Li 2013). Duplicated read alignments were removed using PicardTools MarkDuplicates version 1.95. Variants were called using the Genome Analysis Toolkit (GATK) HaplotypeCaller version 3.7 with default parameters (McKenna *et al*. 2010). Variants were filtered by removing clusters of variants (5 or more variants within 20 bases) using the GATK VariantFiltration tool with parameters “--clusterSize 5 --clusterWindowSize 20”. Variants were subsequently filtered for genotype quality (GQ) < 20 and depth of coverage (DP) < 10 using GATK VariantFiltration with parameters “--genotypeFilterExpression “GQ < 20” --genotypeFilterName GQFilter --genotypeFilterExpression “DP < 10” --genotypeFilterName DPFilter”.

For SNP trees, variants were called using the GATK HaplotypeCaller version 3.7 (McKenna *et al*. 2010) with the additional parameter “--emitRefConfidence GVCF” to produce GVCF files. Joint genotyping was performed for GVCF files from 42 *C. metapsilosis* strains (Table S1) using the GATK GenotypeGVCFs tool with default parameters. SNPs were extracted from the multi-sample VCF and filtered as described above. Repeated Random Haplotype Sampling (RRHS) was used to randomly choose an allele at all heterozygous variant sites and generate a FASTA sequence of all SNPs for each sample (Lischer *et al*. 2014). This process was completed 1000 times to capture the full breadth of allelic variation in the isolates.

Phylogenetic trees were constructed with RAxML version 8.2.9 with the GTRGAMMA model for each of the 1000 SNP sets (Stamatakis 2014). The tree with the best maximum likelihood score was selected as the reference tree, and the remaining 999 trees were used as pseudo-bootstrap trees to generate a supertree.

### Loss of Heterozygosity

Heterozygous regions were defined as regions containing at least two heterozygous variants within 100 base pairs (bp) of each other (Pryszcz *et al*. 2015). Other regions were designated as homozygous. LOH in *C. metapsilosis* MSK414 was further annotated by aligning Illumina reads to the contigs assigned to the A parent from the Canu assembly with BWA-MEM and calling variants with GATK with parameters as described in ‘Variant calling and filtering’ (McKenna *et al*. 2010). LOH regions were annotated as originating from the C parent if they contained at least one homozygous variant, and as originating from the A parent if there were no homozygous variants in the region. LOH blocks were plotted using the karyoploteR package in R (Gel and Serra 2017). Divergence between the haplotypes of *C. metapsilosis* MSK414 was calculated as the number of heterozygous variant sites divided by the total length of the heterozygous regions of the genome.

### Circos plots

To compare the haplotypes of the diploid *C. metapsilosis* MSK414 assembly, contigs were first assigned to haplotypes A and C using BLASTN. Circoletto (with Circos version 0.69) (Darzentas 2010). was used to align the 9 largest contigs assigned to haplotype A to the 9 largest contigs from haplotype C to generate a Circos plot (Krzywinski *et al*. 2009), with options “--e_value 1e-180 --gep 3 --max_ribbons 10000 --hide_orient_lights --z_by alnlen --untangling_off”. To compare the assembly of *C. metapsilosis* ATCC 96143 to the diploid C. metapsilosis MSK414 assembly, the 8 largest contigs from the *C. metapsilosis* ATCC 96143 reference were aligned to the 18 largest contigs from *C. metapsilosis* MSK414 (including haplotypes A and C), with Circoletto. The same procedure was used to compare the 10 largest contigs from the *C. metapsilosis* chimeric reference assembly to the 18 largest contigs from *C. metapsilosis* MSK414 (including haplotypes A and C).

## RESULTS

### Population study of *C. metapsilosis*

The genomes of 11 *C. metapsilosis* isolates were first sequenced in 2015 (Pryszcz *et al*. 2015). All were hypothesized to originate from mating between two related, but genetically distinct, individuals. The two parents differed from each other by ∼4.5% at the genome level. A haploid chimeric reference assembly that comprised 57 contigs totaling 13.4 Mb was constructed by combining data from two strains (Pryszcz *et al*. 2015). Subsequently a collapsed haploid assembly was generated from MinION long read data from *C. metapsilosis* 96413 (Oh *et al*. 2019). Here, we carried out a population genomics analysis of 42 *C. metapsilosis* isolates, including 11 from Pryszcz et al. (Pryszcz *et al*. 2015), one from Oh et al (Oh *et al*. 2019) and 30 from Memorial Sloan Kettering Cancer Center (MSK), of which 25 were described previously (Zhai *et al*. 2020) (Table S1). Most of the MSK strains were collected as part of a study of a cohort of adult patients with culture-proven fungal bloodstream infections following allogeneic hematopoietic stem cell transplant (allo-HCT). Among these, 26 *C. metapsilosis* isolates were isolated from a single patient (Zhai *et al*. 2020). Four were isolated from two other patients with different cancers. DNA was sequenced on an Illumina HiSeq to at least 70X coverage.

The phylogenetic relationship of the *C. metapsilosis* isolates was determined by constructing trees using SNPs identified across all 42 isolates relative to the chimeric reference genome constructed by Pryszcz et al (Pryszcz *et al*. 2015). For heterozygous variant sites, one allele was chosen at random using Repeated Random Haplotype Sampling (RRHS) (Lischer *et al*. 2014). At homozygous variant sites, the alternative allele to the reference was chosen by default. All variant sites were concatenated and SNP trees were drawn using RAxML (Stamatakis 2014). All isolates have high levels of heterozygosity, ranging from 1 heterozygous variant (i.e. heterozygous SNP or indel) per 34 bp to 1 per 49 bp, with an average of 1 per 41 bp (Figure 1B). *C. metapsilosis* MSK414 is distantly related to all other *C. metapsilosis* isolates (Figure 1A). It is also the most heterozygous isolate analyzed, with 398,389 heterozygous variants (1 every 34 bases) (Figure 1B). *C. metapsilosis* MSK414 also has a high number of homozygous variants compared to the *C. metapsilosis* chimeric reference assembly (Pryszcz *et al*. 2015). On average, the *C. metapsilosis* isolates have 53,799 homozygous variants, whereas *C. metapsilosis* MSK414 has 147,375 homozygous variants (Figure 1B).

**Figure 1.**
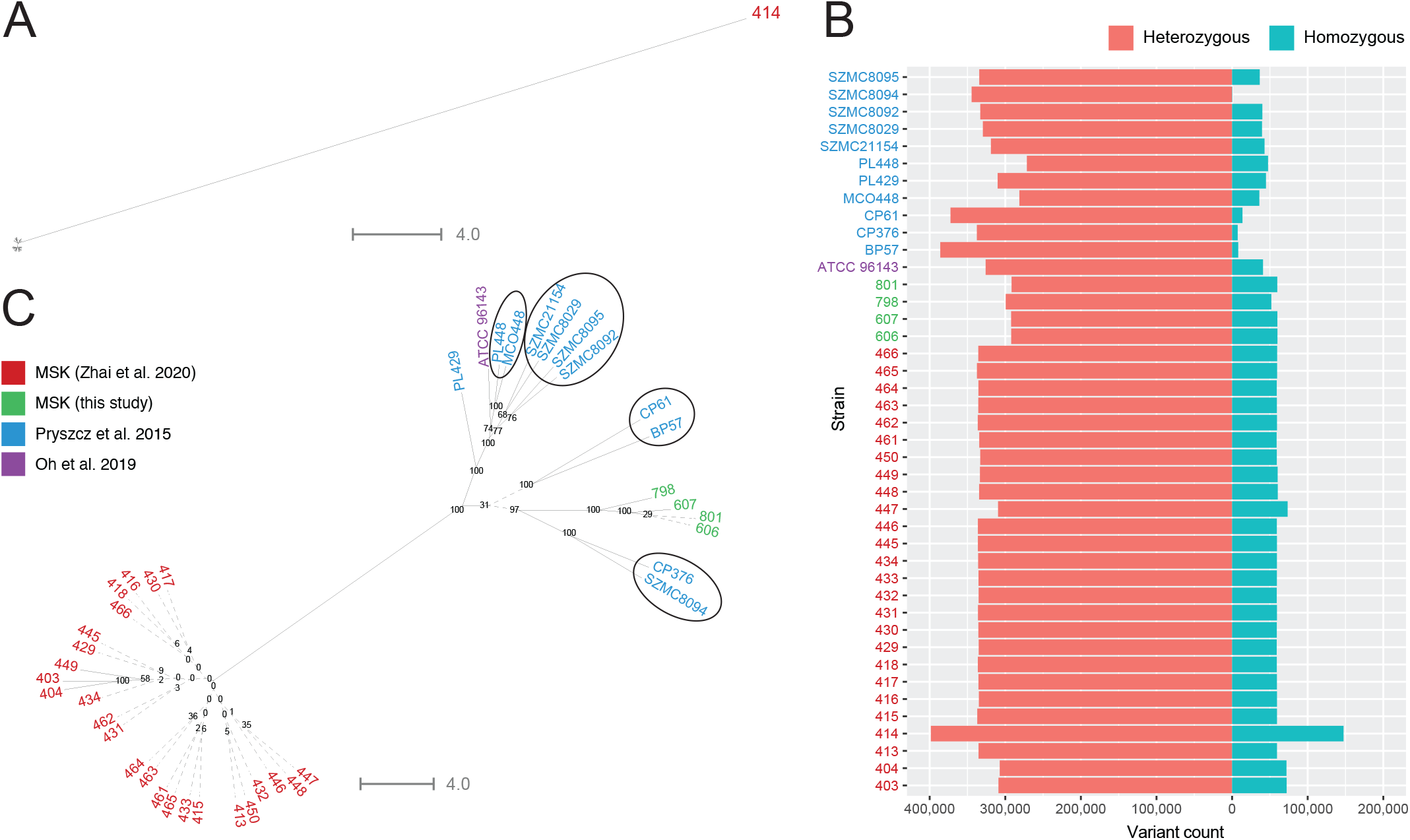
Identification of a divergent *C. metapsilosis* isolate. **A. C. *metapsilosis* MSK414 is highly divergen**t. Phylogenetic SNP trees were generated for 42 clinical *C. metapsilosis* isolates from various geographical regions (Table S1). SNPs were called using GATK HaplotypeCaller and filtered to remove clusters of variants (5 or more variants within 20 bases) and variants with genotype quality (GQ) < 20 or depth of coverage (DP) < 10 using the GATK Variant-Filtration tool. Repeated Random Haplotype Sampling (RRHS) was used to randomly choose an allele at all heterozygous variant sites and generate a FASTA sequence of all SNPs for each sample (Lischer et al., 2014). In the case of homozygous SNPs, the alternate allele was chosen by default. This process was repeated 1000 times and 1000 phylogenetic trees were constructed with RAxML using the GTRGAMMA model (Stamatakis, 2014). The tree with the best maximum likelihood score was selected as the reference tree, and the remaining 999 trees were used as pseudo-bootstrap trees to generate a supertree. Pseudo-bootstrap values are shown as branch labels. *C. metapsilosis* MSK414 is labeled in red, while other isolates are not labeled. **B. *C. metapsilosis* MSK414 has more variants than any other *C. metapsilosis* isolate**. Variant count is shown on the bidirectional X-axis, with heterozygous variants shown on the left in orange and homozygous variants shown on the right in blue. *C. metapsilosis* strains are labelled on the Y-axis. Isolates from MSK are labelled without the “MSK” prefix. Heterozygosity levels range from 271,440 to 398,389 heterozygous variants. *C. metapsilosis* MSK414 has more heterozygous variants than all other isolates. Some isolates have almost no homozygous variants, e.g. *C. metapsilosis* isolates SZMC8094 (used to construct the reference assembly), CP61, CP376 and BP57. *C. metapsilosis* MSK414 has more heterozygous variants and more than double the number of homozygous variants of any other *C. metapsilosis* isolate. **C. Other *C. metapsilosis* isolates fall into two main clusters**. Phylogenetic trees for all *C. metapsi-losis* isolates except MSK414 were drawn as in part A. Isolates from MSK are labelled without the “MSK” prefix. Isolates described by Zhai et al. (Zhai et al. 2020) cluster together and are highly similar. Their relationships cannot be accurately resolved (indicated by dashed lines, bootstrap < 40). Four MSK isolates, *C. metapsilosis* MSK606, *C. metapsilosis* MSK607, *C. metapsilosis* MSK798 and *C. metapsilosis* MSK801, cluster together and are more similar to the clinical isolates described by Pryszcz et al. (Pryszcz et al. 2015) and Oh et al. (Oh et al. 2019) than the other isolates from MSK. The inferred phylogenetic relationships of the isolates analysed by Pryszcz et al. (Pryszcz et al. 2015) fall into four groups, supporting the original analysis, represented by black circles.

To facilitate a comparison among the other *C. metapsilosis* isolates, SNP trees were drawn excluding *C. metapsilosis* MSK414 (Figure 1C). 25 of the 26 *C. metapsilosis* MSK strains (designated by four as the first digit) isolated from a single patient cluster together, as described previously (Zhai *et al*. 2020). The genomes of these isolates are highly similar (Figure 1C) and could not be differentiated by phylogenetic analysis, although there are some differences in homozygosity levels (Figure S1). Four additional *C. metapsilosis* isolates from MSK (labeled in green on Figure 1C) cluster separately from the other MSK strains. Isolates described by Pryszcz et al. (Pryszcz *et al*. 2015) fall into approximately four clades (encircled in Figure 1C), as previously described. *C. metapsilosis* PL429 does not belong to any clade. *C. metapsilosis* ATCC 96143, a clinical isolate from Livermore, USA, clusters with one of the groups previously identified by Pryszcz et al. (Pryszcz *et al*. 2015), together with *C. metapsilosis* MCO448 and *C. metapsilosis* PL448, which are both clinical isolates from Washington, USA.

### Identification of a novel *C. metapsilosis* hybrid

Figure 1A shows that *C. metapsilosis* MSK414 is very different to the other *C. metapsilosis* isolates. We therefore attempted to assemble its genome to facilitate comparison. Previous studies have shown that there are many limitations associated with assembly of short read data from heterozygous diploids (Chan *et al*. 2012; Zheng *et al*. 2013; Pryszcz and Gabaldón 2016). During assembly of most diploid genomes, the two haplotypes collapse into a single contig, yielding a haploid assembly. However, for highly heterozygous genomes, this is not possible, and the resulting assemblies are highly fragmented (Pevzner *et al*. 2001; Li *et al*. 2010; Gnerre *et al*. 2011). Pryszcz and Gabaldón (Pryszcz and Gabaldón 2016) developed a protocol (Redundans) that produces a haploid reference assembly by collapsing sequence information from both haplotypes. At heterozygous sites, one allele is randomly chosen to generate one representative contig per diploid chromosome. They assembled a chimeric *C. metapsilosis* haploid genome, using data from two isolates, that has 57 contigs (Pryszcz *et al*. 2015). However, haplotype information has been lost from this assembly.

We used SPAdes (Bankevich *et al*. 2012), which keeps haplotypes separate, to assemble the genomes of 42 *C. metapsilosis* isolates (Table S1). Scaffolds fell into two groups, where the depth of coverage of one group was approximately half of the coverage of the second group. Scaffolds with half coverage represent heterozygous regions where both haplotypes have been assembled separately. Scaffolds with high depth of coverage represent homozygous regions that have been collapsed into a single scaffold. This is shown for *C. metapsilosis* MSK414 (Figure 2) using a coverage-versus-length (CVL) plot (Douglass *et al*. 2019). This assembly pattern suggests that like all other *C. metapsilosis* isolates, *C. metapsilosis* MSK414 is a hybrid.

**Figure 2.**
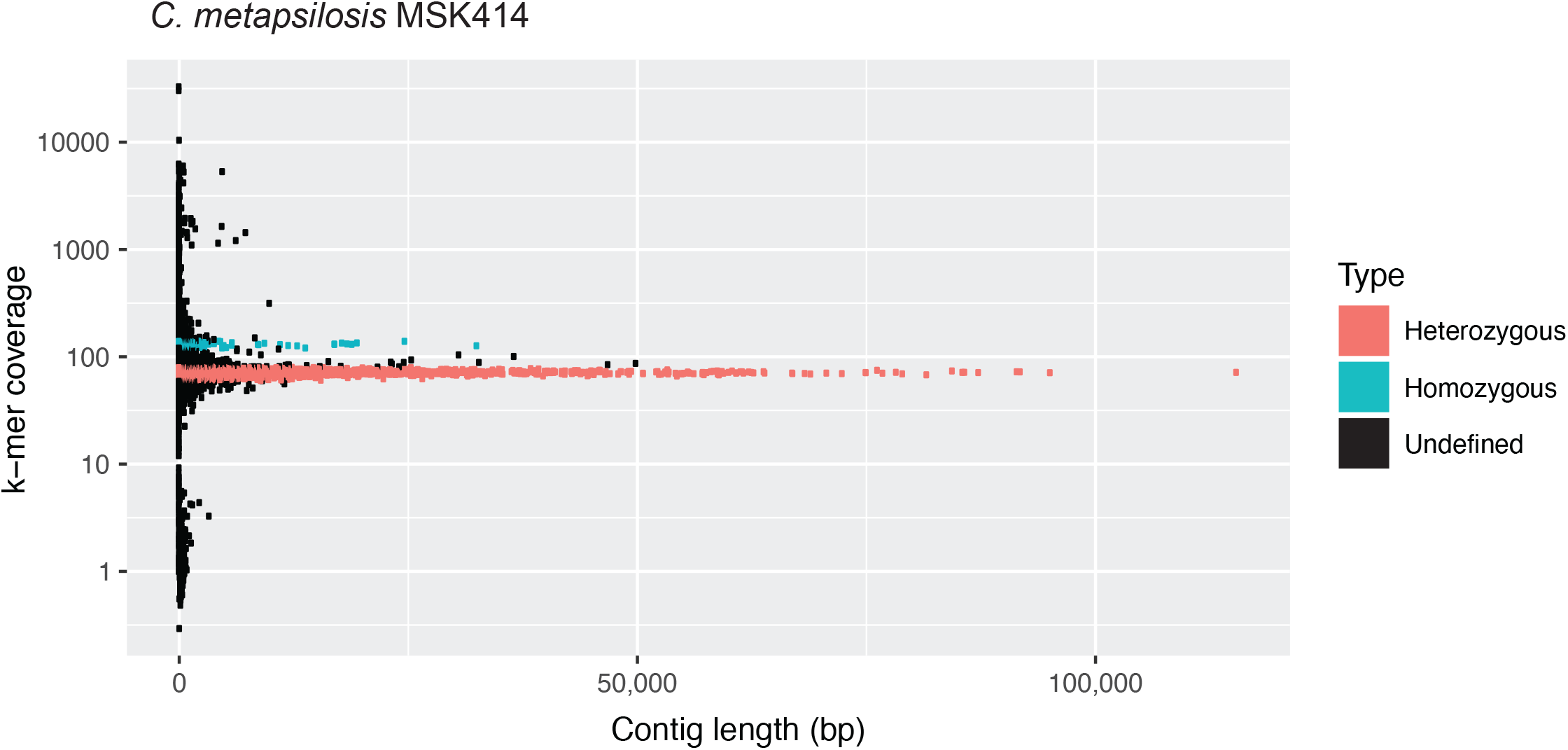
Illumina assembly of *C. metapsilosis* MSK414 reveals two peaks of coverage. Scaffolds from the SPAdes assembly of the Illumina data from *C. metapsilosis* MSK414 are shown as dots. Scaffold length is shown on the X-axis and scaffold k-mer coverage is shown on the Y-axis on a log scale. The majority of the scaffolds have approximately 70X coverage (red). These scaffolds represent heterozygous regions, where both haplotypes have been assembled separately. A second peak of coverage is visible at approximately 130X (cyan). These scaffolds represent homozygous regions that have been collapsed. This structure suggests that *C. metapsilosis* MSK414 has a hybrid genome (i.e. the two haplotypes are distinctly different).

To improve the assembly of *C. metapsilosis* MSK414, we used Oxford Nanopore MinION long read sequencing. The reads were assembled using Canu (Koren *et al*. 2017), and errors were corrected by incorporating the Illumina data using Pilon (Walker *et al*. 2014). This generated an assembly of 45 contigs, with 18 larger than 450 kb, totaling 27 Mb (Table S2). The contigs smaller than 450 kb were derived from the mitochondrial genome, or from within the chromosomal-sized contigs. A telomeric repeat (ACTTTGGACATCCTAACCTCAAT) was identified at both ends of 14 contigs, and at one end of three of the largest contigs in the assembly. Centromeres (Ola *et al*. 2020) were identified in 16 contigs, which is consistent with hybridization between two parents with eight chromosomes each (Table S3).

To identify the two haplotypes, we compared the contigs to each other (Figure 3). There is a direct relationship between 13 of the 18 largest contigs. Based on similarities (Figure 3A and Table S3) we assume that tig11866 and tig3 should be joined, and they represent the haplotype A equivalent of tig1 from haplotype C (Figure 3). These 13 contigs therefore represent both haplotypes for 6 of the 8 pairs of *C. metapsilosis* chromosomes. However, the remaining two chromosome pairs are not collinear. tig10 from one haplotype (haplotype A) matches parts of both tig11870 and tig11878 from the second haplotype (haplotype C) (Figure 3A). Similarly, tig11881 from one haplotype (haplotype A) matches part of tig11878 and tig11874 from the second haplotype (haplotype C). Based on similarities, we assume that tig11870 and tig11874 (haplotype C) should be joined (Figure 3B). This is consistent with a single translocation event between the two parental haplotypes (Figure 3B). The translocated chromosomes contain the mating-type like loci (MTL).

**Figure 3.**
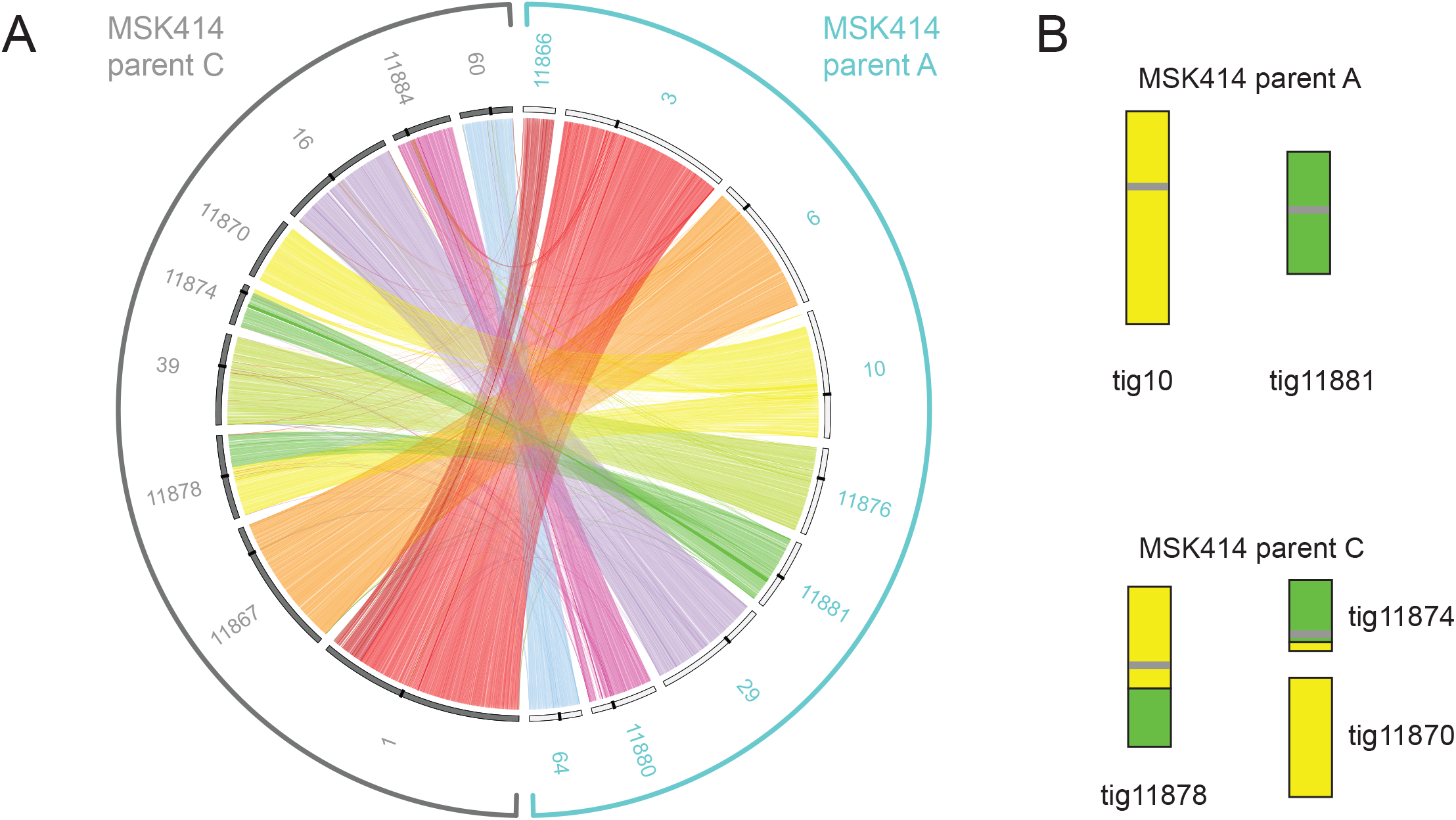
Haplotypes A and C in *C. metapsilosis* MSK414 differ by one reciprocal translocation. A. Similarity between the haplotypes of *C. metapsilosis* MSK414 was visualized using Circos (Krzywinski et al. 2009) and Circoletto (Darzentas 2010). The 18 largest contigs in the assembly are shown, with the haplotype from the putative A parent on the right (outlined in turquoise) and from the putative C parent on the left (outlined in gray). For clarity, contigs are labelled without the “tig” prefix. Sequences with similarity were identified by BLASTN and alignments with a minimum E-value (1e-180) were plotted as links between the two haplotypes. The 9 largest contigs from parent A (shared with other *C. metapsilosis* isolates) are shown on the right hand side with white bars on the inner layer. The 9 largest contigs from parent C are shown on the left with gray bars on the inner layer. Centromeres are shown as black bars on the inner layer. A translocation is evident between tig10 and tig11881 in the A haplotype and the equivalent contigs in the C haplotype. B. Translocation between tig10 and tig11881 from haplotype A in haplotype C of *C. metapsilosis* MSK414. Contigs in the *C. metapsilosis* parent A and C haplotypes are shown as colored bars. Centromeres are shown as gray horizontal bars on the contigs.

To assign the contigs to haplotypes, we compared them to a haploid assembly of *C. metapsilosis* ATCC 96143 (Oh *et al*. 2019). This assembly is more complete (8 scaffolds) than the original chimeric reference assembly generated by Pryszcz et al (Pryszcz *et al*. 2015), but still represents a collapsed haploid. In most cases, there is a 1:2 relationship between the haploid assembly and the *C. metapsilosis* MSK414 contigs (Table S3, Figure 3). For each of these, one *C. metapsilosis* MSK414 contig is more similar to the reference (94-96% identity) and one is less similar (92-93%). These likely represent the haplotypes of the original parents of MSK414. Contigs 3.1 and 5.1 of *C. metapsilosis* ATCC 96143 match two contigs in one haplotype of *C. metapsilosis* MSK414 because of the reciprocal translocation (Figure 3B).

We assigned the set of contigs that are more similar to *C. metapsilosis* ATCC 96143 as haplotype A, and the set of contigs that are less similar to C. *metapsilosis* ATCC 96143 as haplotype C (Table S3).

### Analysis of the Mating-type Like Locus

Pryszcz et al. (Pryszcz *et al*. 2015) showed that *MTLalpha* is intact in 11 *C. metapsilosis* isolates, and is identical in the order and orientation of its genes to the *MTLalpha* locus in *C. albicans, C. tropicalis* and *C. orthopsilosis*. The *MTL***a** locus, however, has been partially overwritten with information from the *MTLalpha* locus (Figure 4). In *MTL***a**, the *PAP***a** and *OBP***a** genes have been replaced by the *MTLalpha*2 and *OBPalpha* genes. A portion of *PIK***a** has been overwritten with a portion of *PIKalpha*, producing a chimeric *PIK* gene at the *MTL***a**. We found the same *MTL* arrangement in 30 additional *C. metapsilosis* isolates (*C. metapsilosis* ATCC 96143 and 29 *C. metapsilosis* isolates from MSK). However, *C. metapsilosis* MSK414 has a different organization; both *MTL***a** and *MTLalpha* are intact (Figure 4).

**Figure 4.**
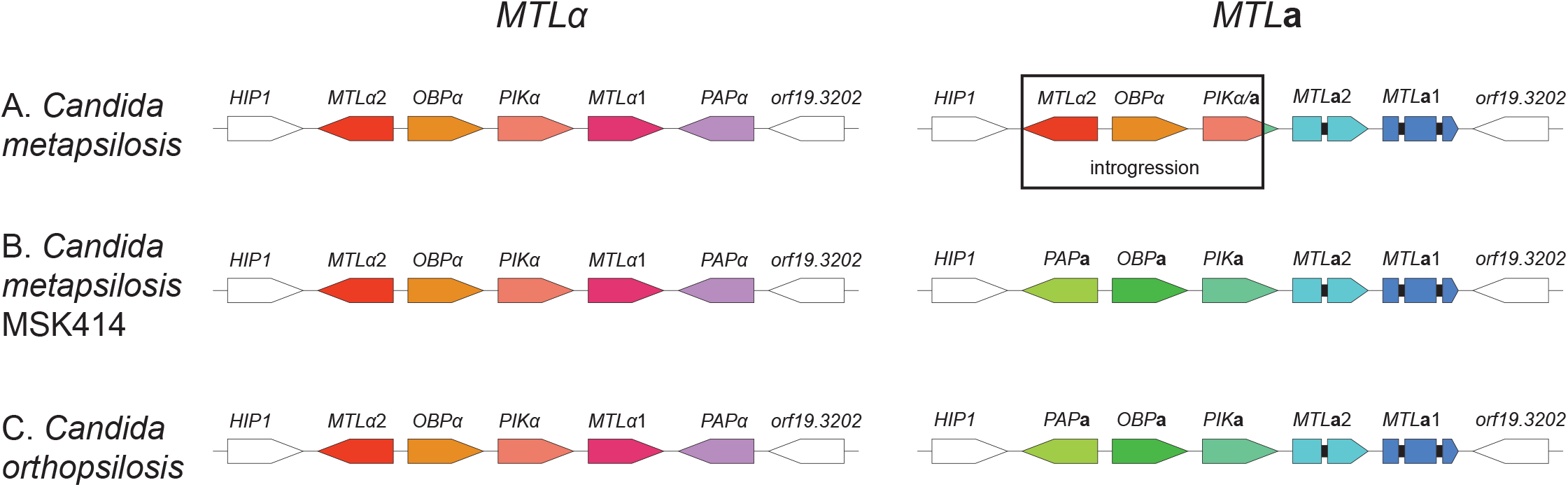
*C. metapsilosis* MSK414 has a distinct arrangement at MTLa. The structure of the MTL**a** and MTLalpha idiomorphs in the majority of *C. metapsilosis* isolates (A), *C. metapsilosis* MSK414 (B), and *C. orthopsilosis* (C) are shown, with their relative orienta-tions. Introns in the MTL**a**1 and MTL**a**2 genes are shown as black bars. In most *C. metapsilosis* isolates, the MTLalpha locus is intact, and is identical in its order and orientation to that of *C. orthopsilosis*. The MTL**a** locus has been partially overwritten by a portion of the MTLalpha locus containing the genes MTLalpha2, OBPalpha, and a part of PIKalpha. The PIK gene is chimeric, comprising part of PIKalpha and part of PIK**a**. Both MTL idiomorphs in *C. metapsilosis* isolate MSK414 (B) have the same order and orientation as in *C. orthopsilosis* (C). In this isolate, the MTLalpha and MTLa idiomorphs are intact with no introgression.

The *MTLalpha* locus from *C. metapsilosis* MSK414 is ∼99.8% identical to the *MTLalpha* locus from *C. metapsilosis* ATCC 96143. In addition, the copy of *orf19*.*3202* that is adjacent to *MTLalpha* is 97% identical to the reference genome, whereas the copy adjacent to *MTL***a** is only 92% identical. We therefore assume that *MTLalpha* was contributed by the same parent, or a very similar parent, in all previously described *C. metapsilosis* isolates and in *C. metapsilosis* MSK414 (parent A). For most *C. metapsilosis* isolates, a second parent (parent B) donated the *MTL***a** locus, which has subsequently been overwritten. In *C. metapsilosis* MSK414 however, *MTL***a** was donated by a third parent, parent C. The majority of *C. metapsilosis* isolates are AB hybrids, whereas *C. metapsilosis* MSK414 is an AC hybrid.

### Loss of heterozygosity

Previous studies observed that *C. metapsilosis* isolates have undergone large-scale LOH events (Pryszcz *et al*. 2015). In addition, we found that *C. metapsilosis* isolates MSK403, MSK404 and MSK447 have undergone LOH across most of scaffold 4 (Figure S1). All isolates except for *C. metapsilosis* MSK414 have undergone significant LOH across part of two scaffolds (Figure S1), supporting the hypothesis that they all descended from the same ancestor. Each *C. metapsilosis* genome has undergone LOH over approximately half its length (Figure 5A).

**Figure 5.**
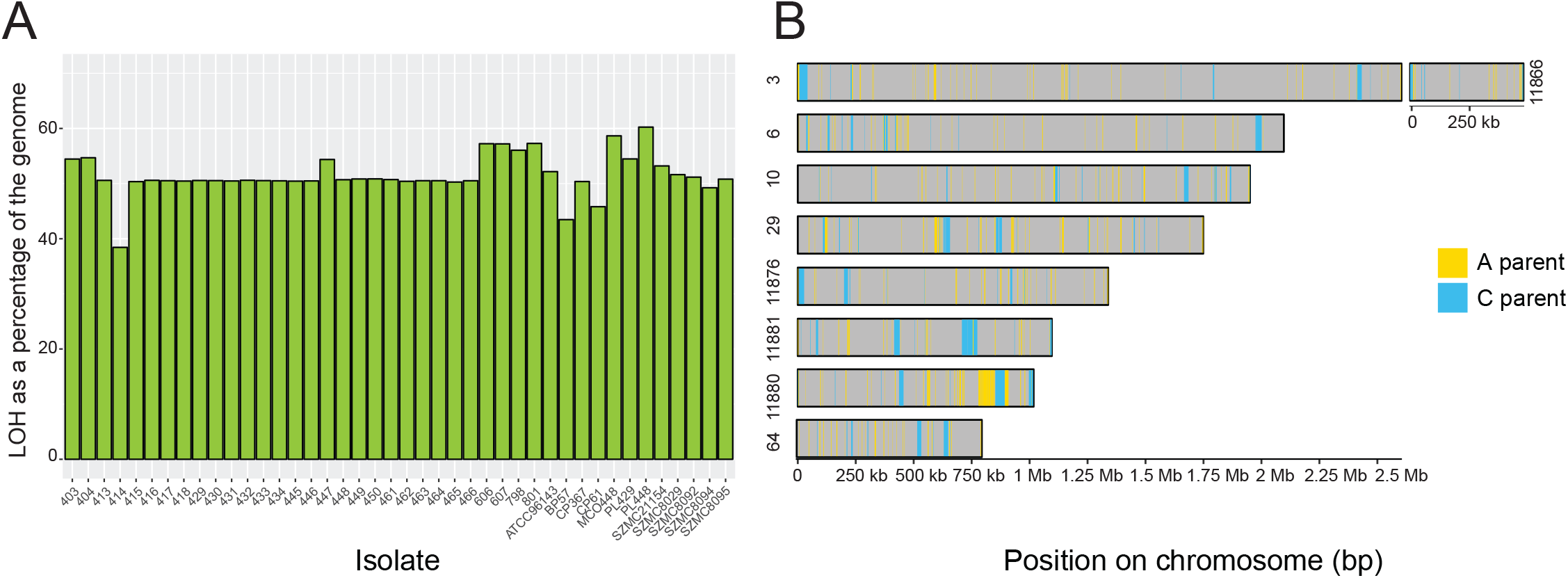
The genome of *C. metapsilosis* MSK414 has undergone less LOH than other *C. metapsilosis* isolates. A. Percentage of the genome that has undergone LOH (Y-axis) in *C. metapsilosis* isolates (X-ax-is). For most isolates, more than 50% of the genome has undergone LOH, equating to approxi-mately 6.7 Mb. Only 38% of the *C. metapsilosis* MSK414 genome has undergone LOH. Two other isolates, *C. metapsilosis* BP57 and *C. metapsilosis* CP61, which are closely related (Fig. 1), have also undergone less LOH than the other isolates (approximately 43 and 45% respec-tively). B. Regions of LOH in *C. metapsilosis* MSK414 are dispersed throughout the genome. The nine largest contigs assigned to the *C. metapsilosis* MSK414 A haplotype are shown. For the sake of clarity, only LOH regions of at least 1 kb are illustrated here. Heterozygous regions (defined as any region with at least 2 heterozygous variants within 100 bp of each other), undefined regions, and LOH regions less than 1 kb are colored in gray. LOH blocks were defined as any region of at least 100 bp with fewer than 2 heterozygous variants. LOH regions were assigned to the A parent haplotype (colored in yellow) if there were any homozygous variants present, and to the C parent haplotype (colored in blue) if there were no homozygous variants.

The novel hybrid MSK414 stands out as having undergone relatively little LOH (38% of its length; 5.2 Mb) (Figure 5A). Regions of LOH were assigned to either the C parent (at least one homozygous variant in 100 bp) or to the A parent (no homozygous variants). LOH regions assigned to parent C totaled ∼5.4% of the total genome length, while LOH assigned to the A parent totaled ∼ 26% of the total genome length. There are 350 blocks of LOH with an average length of 251 bp assigned to the C haplotype, and 13,892 blocks with an average length of 2,080 bp assigned to the A haplotype. The LOH tracts are randomly dispersed throughout the contigs (Figure 5B).

## DISCUSSION

Previously sequenced *C. metapsilosis* isolates have an intact *MTLalpha* locus, with introgression at *MTL***a** (Pryszcz *et al*. 2015). We found this arrangement in 29 additional isolates from the USA (MSK) and one from Italy (*C. metapsilosis* ATCC 96143). The relative lack of divergence among these isolates (Figure 1B), and the observation that most share LOH tracts, suggest that they are all derived from the same hybrid ancestor. It is likely that a single ancient hybridization event between A and B parents that differ by 4.5% was followed by introgression at *MTL***a**, and that most *C. metapsilosis* isolates descended from this single event.

Despite a lack of evidence at the time, Pryszcz et al. (Pryszcz *et al*. 2015) suggested that additional hybrid lineages of *C. metapsilosis* may be found. Indeed, analysis of other fungal species, including *Cryptococcus neoformans* and *C. orthopsilosis*, showed that hybridization in those species is ongoing and has occurred on multiple separate occasions (Xu *et al*. 2002; Li *et al*. 2012; Schröder *et al*. 2016). However, until now, no different hybrids of *C. metapsilosis* have been identified. Our results show that *C. metapsilosis* MSK414 most likely shares one parent (A) with other *C. metapsilosis* isolates, but its second parent (C) is distinctly different. Parent C has donated an intact *MTL***a** idiomorph. The A and C haplotypes differ by approximately 4.46%, similar to the divergence between the A and B haplotypes in the other *C. metapsilosis* isolates (4.5%; (Pryszcz *et al*. 2015)). This is also similar to the divergence between haplotypes in hybrids of *C. orthopsilosis* (Schröder *et al*. 2016) and *C. tropicalis* (O’Brien *et al*. 2021).

Identification and separation of parental haplotypes in hybrid species is difficult unless at least one of the parents is known. For *C. orthopsilosis*, the first genome sequence came, fortuitously, from a highly homozygous strain (*C. orthopsilosis* 90-125) and so provided a pure reference sequence for the A haplotype of this species (Riccombeni *et al*. 2012). Subsequent studies revealed that the majority of *C. orthopsilosis* isolates are, in fact, hybrids between one parent that is essentially identical to this homozygous reference strain, and a second parent that is approximately 4.5% different from it (Pryszcz *et al*. 2014; Schröder *et al*. 2016). For *C. metapsilosis* MSK, we were able to separate the haplotypes using long read sequencing (ONT).

Pure lineages of the A, B and C haplotypes of *C. metapsilosis* have not yet been identified. Pryszcz et al (Pryszcz *et al*. 2015) suggested that only hybrid lineages of *Candida* species are pathogenic, and that homozygous isolates may only be found in non-clinical samples. This proposal is supported by the observation that most clinical isolates of *C. orthopsilosis* are hybrids (Schröder *et al*. 2016), and that *C. albicans* is an ancient hybrid (Mixão and Gabaldón 2020). However, rare hybrids of *C. tropicalis* are enriched in environmental, and not clinical sites (O’Brien *et al*. 2021). Although *C. metapsilosis* has been isolated from several different body sites, including blood, feces, mucosa, nails, skin, and urine (Hensgens *et al*. 2009), its natural environment is not known. Hybridization may have enabled *C. metapsilosis* to colonies a new niche, namely the human host. However, “AC” hybrids are rare (1 of 42 isolates), and the effect on pathogenicity cannot be fully characterised until environmental isolates are identified.

Our results, together with studies in other species such as *C. albicans* (Mixão and Gabaldón 2020), *C. orthopsilosis* (Pryszcz *et al*. 2014; Schröder *et al*. 2016), *C. tropicalis* (O’Brien *et al*. 2021) and *Millerozyma sorbitophila* (Louis *et al*. 2012), suggest that hybridization occurs frequently in members of the CUG-Ser1 clade, and is likely to be current and ongoing. Hybridization may represent a mode of adaptation to the host, or possibly to other as yet undetermined conditions.

## Data availability

All strains are available by request. The raw Illumina data for *C. metapsilosis* MSK606, MSK607, MSK798, MSK801 and MSK414 are available at accession numbers XXXXX (awaiting accession number). The raw MinION data for *C. metapsilosis* MSK414 is available at accession number SRR15054248, and the genome assembly is available at BioProject PRJNA730502 (Accession number JAHFZM000000000).

## Data availability

The raw Illumina data for *C. metapsilosis* MSK606, MSK607, MSK798, MSK801 and MSK414 are available at accession numbers XXXXX (awaiting accession number). The raw MinION data for *C. metapsilosis* MSK414 is available at accession number SRR15054248, and the genome assembly is available at BioProject PRJNA730502 (Accession number JAHFZM000000000).

## Acknowledgements

This work was supported by the Wellcome Trust (grant number 109167/Z/15/Z) (G.B.), and Science Foundation Ireland (grant number 19/FFP/6668) (G.B.), Deutsche Forschungsgemeinschaft (DFG, German Research Foundation) grant RO-5328/2 (T.R.), National Institutes of Health (NIH) grants R01 AI093808 (T.M.H.), R21 AI156157 (T.M.H.), P30 CA008748 (Cancer Center Core Grant), the Ludwig Center for Cancer Immunotherapy (T.M.H.), and the Susan and Peter Solomon Divisional Genomics Program (T.M.H.). For the purpose of Open Access, the authors have applied a CC BY public copyright license to any Author Accepted Manuscript version arising from this submission

---

**Supplementary Figure 1.**
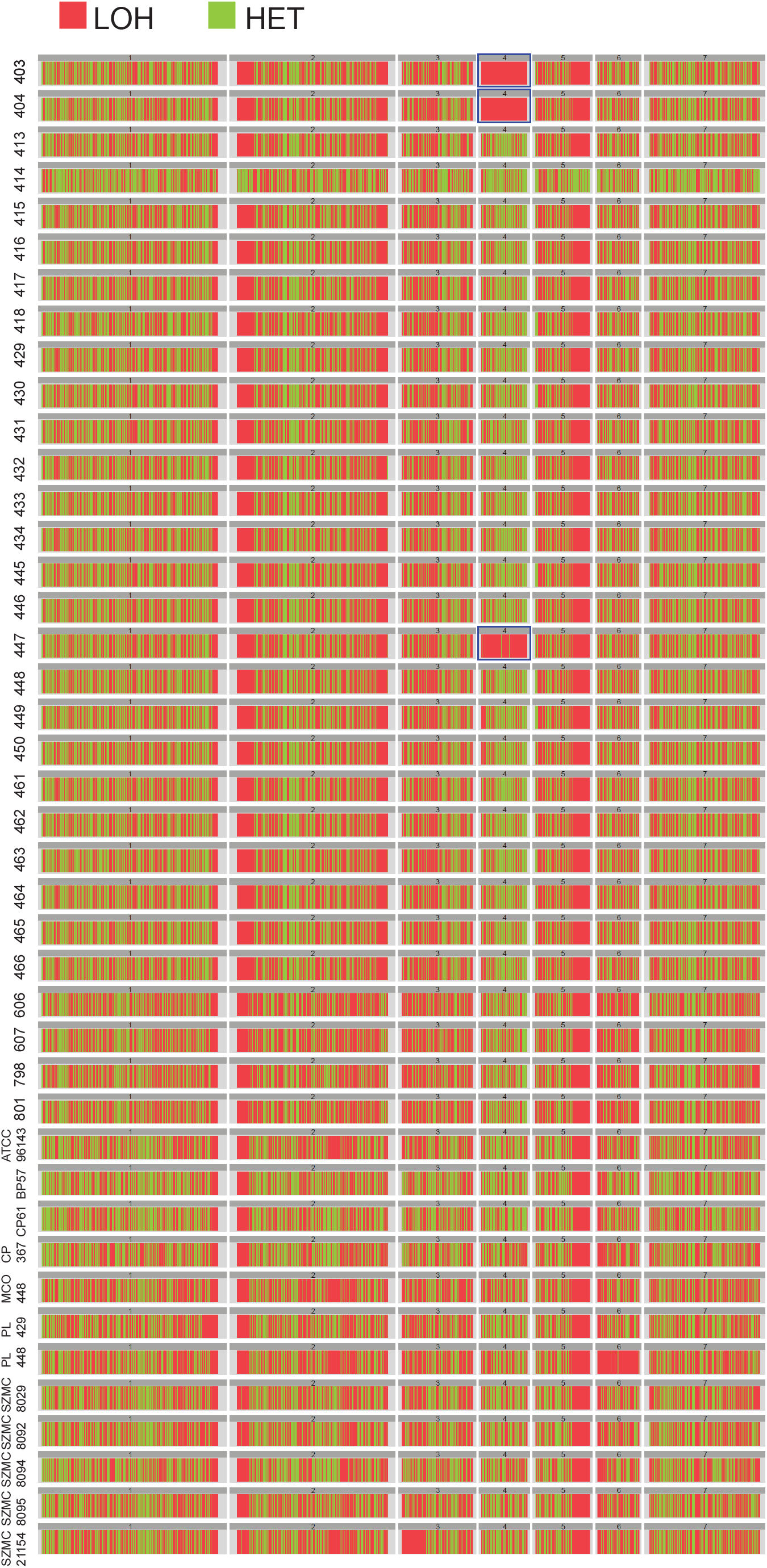
Distri-bution of heterozygous and LOH regions in the genomes of *metapsilosis* isolates. The seven largest scaffolds in the chimeric reference genome are displayed horizontally from left to right and labelled from 1 to 7. Regions of LOH are shown in red and heterozygous (HET) regions are shown in green. Isolates are labelled on the left-hand side. MSK isolates are shown without the “MSK” prefix. The genomes of most isolates consist of a mixture of heterozygous and LOH regions. Isolates 403, 404, and 447 have undergone significant LOH across most of scaffold 4 (highlighted with blue boxes). Isolate PL448 has undergone significant LOH on scaffold 6. Large areas of LOH are visible on scaffolds 2 and 5 in all isolates, except for the new hybrid *C. metapsilosis* MSK414.

**Supplementary Figure 2.**
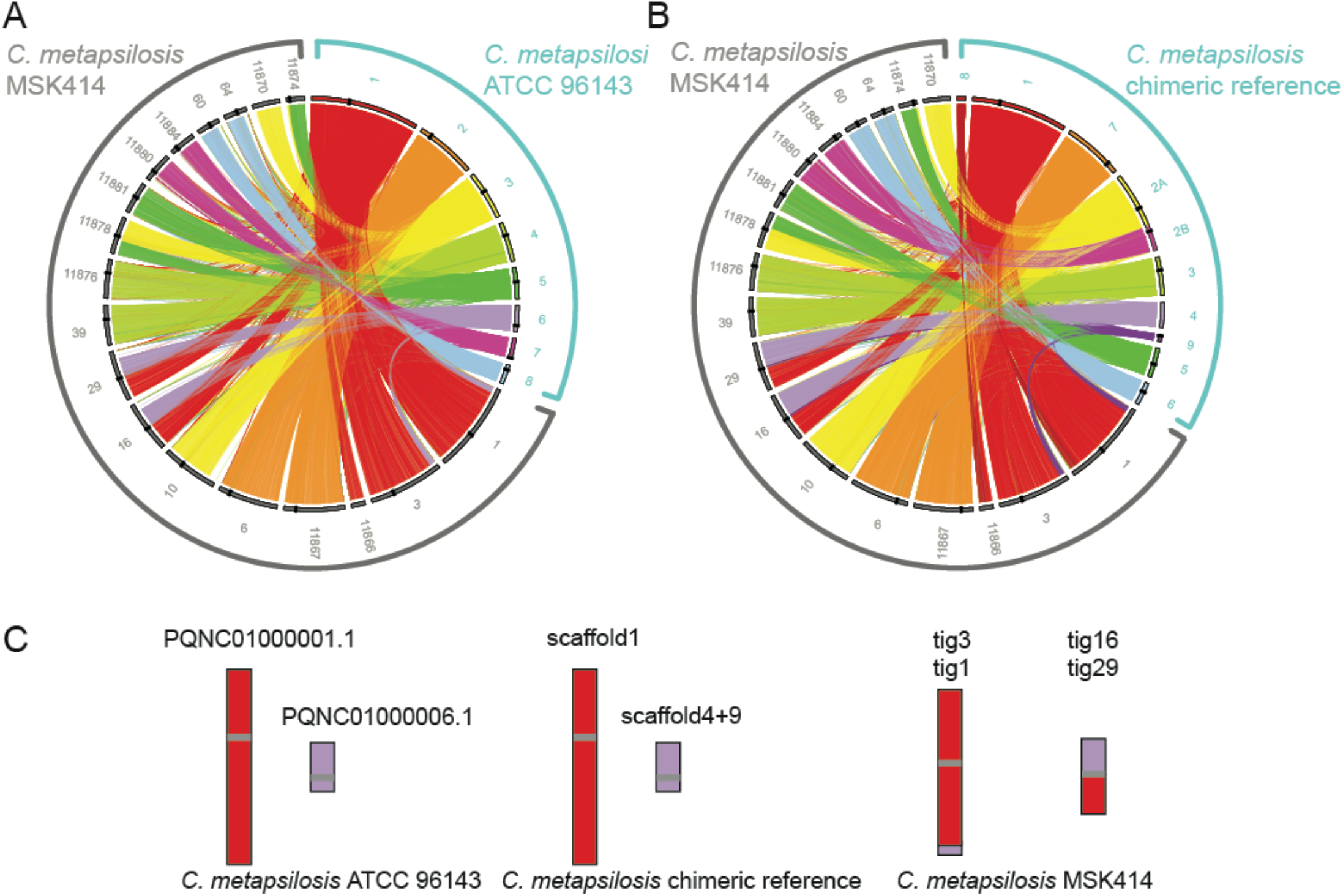
Rearrangements between the diploid assembly of *C. metapsilosis* MSK414 and two haploid *C. metapsilosis* assemblies. Sequence similarity between the genome assembly of *C. metapsilosis* MSK414 and the assembly of *C. metapsilosis* ATCC 96143 (A) and the chimeric reference assembly of *C. metapsilosis* (B) was visualized using Circos (Krzywinski et al. 2009) and Circoletto (Darzentas 2010). Sequences with similarity were identified by BLASTN and alignments with the minimum E-value (1e-180)were plotted as links between the two assemblies. The 18 largest contigs from the diploid *C. metapsilosis* MSK414 assembly are represented by grey bars and the “tig” prefix removed for clarity in panels A and B. Centromeres are marked as black bars on the inner layer. A. Rearrangements between the assembly of *C. metapsilosis* MSK414 and the assembly of *C. metapsilosis* ATCC 96143. The eight largest contigs from the haploid *C. metapsilosis* ATCC 96143 assembly are shown on the upper right hand side of the Circos plot in panel A with colored bars and outlined in turquoise. A mismatch between PQNC01000001.1 and PQNC01000006.1 is observed in both haplotypes of *C. metapsilosis* MSK414 (tig1 and tig3, and tig16 and tig29). A translocation in one haplotype of *C. metapsilosis* MSK414 is apparent between PQNC01000003.1 and PQNC01000005.1 (tig11878 and tig11870/tig11874). B. Rearrangements between the assembly of *C. metapsilosis* MSK414 and the *C. metapsilosis* chimeric reference assembly. The nine largest contigs from the haploid *C. metapsilosis* ATCC 96143 assembly are shown on the upper right hand side of the Circos plot in panel B with colored bars and labelled in turquoise. Scaffold 2 contains two centromeres and appears to be the product of an assembly error (supported by the haploid assembly of *C. metapsilosis* ATCC 96143 and tig10 of the *C. metapsilosis* MSK414 assembly). We refer these as scaffold 2A and scaffold 2B. A mismatch between scaffold 1 and scaffold 4/9 (which we assume should be joined) is visible in both haplotypes of *C. metapsilosis* MSK414 (tig1 and tig3, tig16 and tig29). Scaffolds 4 and 9 most likely represent one chromosome but the assembly has broken at the centromere. This rearrangementmirrors that between scaffolds 1 and 6 of the *C. metapsilosisATCC* 96143 assembly. C. Error in the assembly of *C. metapsilosis* ATCC 96143 and the chimeric *C. metapsilosis* reference assembly. Contigs in the *C. metapsilosisATCC* 96143 assembly, the chimeric *C. metapsilosis* reference assembly and the *C. metapsilosis* MSK414 Canu assembly are shown as colored bars. Centromeres are shown as gray horizontal bars on the contigs. The rearrangement observed between PQNC010000001.1 and PQNC010000006.1 is equivalent to the rearrangement between scaffold 1 and scaffold 4/9 in the *C. metapsilosis* chimeric reference. This is likely to be an assembly error in the two haploid assemblies.

**Table S1.** *C. metapsilosis* strains used in this study.

| **Strain ID** | **Origin** | **Site of Isolation** | **Read length** | **Cov** | **Total variants** |
| --- | --- | --- | --- | --- | --- |
| ATCC 96143<sup>1</sup> | Livermore, USA | Unknown | 150 | 42 | 366,857 |
| BP57<sup>2</sup> | Pécs, Hungary | Throat | 96 | 151.31 | 394,081 |
| CP376<sup>2</sup> | Pisa, Italy | Feces | 96 | 163.96 | 345,030 |
| CP61<sup>2</sup> | Pisa, Italy | Nail | 96 | 151.42 | 386,151 |
| MCO448<sup>2</sup> | Washington, USA | Hand | 46 | 175.76 | 317,333 |
| PL429<sup>2</sup> | Livermore, USA | Unknown | 76 | 284.81 | 354,922 |
| PL448<sup>2</sup> | Washington, USA | Hand | 46 | 186.37 | 318,964 |
| SZMC211154<sup>2</sup> | Cataluña, Spain | Blood | 100 | 492.44 | 361,914 |
| SZMC8029<sup>2</sup> | Debrecen, Hungary | Blood | 100 | 425.57 | 369,230 |
| SZMC8092<sup>2</sup> | Pisa, Italy | Lung | 100 | 470.78 | 373,057 |
| SZMC8094<sup>2</sup> | Pisa, Italy | Feces | 100 | 402.91 | 344,840 |
| SZMC8095<sup>2</sup> | Pisa, Italy | Nail | 100 | 480.67 | 370,999 |
| MSK403<sup>3</sup> | New York, USA | Blood | 101 | 205.47 | 381,084 |
| MSK404<sup>3</sup> | New York, USA | Blood | 101 | 248.1 | 379,288 |
| MSK413<sup>3</sup> | New York, USA | Feces | 101 | 262.04 | 394,729 |
| MSK414<sup>4</sup> | New York, USA | Feces | 101 | 258.03 | 545,764 |
| MSK415<sup>3</sup> | New York, USA | Feces | 101 | 226.01 | 396,420 |
| MSK416<sup>3</sup> | New York, USA | Feces | 101 | 244.2 | 394,311 |
| MSK417<sup>3</sup> | New York, USA | Feces | 101 | 285.59 | 394,741 |
| MSK418<sup>3</sup> | New York, USA | Feces | 101 | 267.76 | 395,401 |
| MSK429<sup>3</sup> | New York, USA | Feces | 101 | 314.98 | 394,129 |
| MSK430<sup>3</sup> | New York, USA | Feces | 101 | 293.11 | 394,695 |
| MSK431<sup>3</sup> | New York, USA | Feces | 101 | 213.65 | 395,459 |
| MSK432<sup>3</sup> | New York, USA | Feces | 101 | 226.58 | 394,260 |
| MSK433<sup>3</sup> | New York, USA | Feces | 101 | 257.45 | 394,628 |
| MSK434<sup>3</sup> | New York, USA | Feces | 101 | 277.28 | 395,131 |
| MSK445<sup>3</sup> | New York, USA | Feces | 101 | 297.49 | 395,536 |
| MSK446<sup>3</sup> | New York, USA | Feces | 101 | 314.58 | 395,372 |
| MSK447<sup>3</sup> | New York, USA | Feces | 101 | 275.69 | 383,033 |
| MSK448<sup>3</sup> | New York, USA | Feces | 101 | 256.41 | 395,011 |
| MSK449<sup>3</sup> | New York, USA | Feces | 101 | 284 | 393,777 |
| MSK450<sup>3</sup> | New York, USA | Feces | 101 | 234.69 | 392,162 |
| MSK461<sup>3</sup> | New York, USA | Feces | 101 | 257.75 | 393,295 |
| MSK462<sup>3</sup> | New York, USA | Feces | 101 | 284.83 | 395,788 |
| MSK463<sup>3</sup> | New York, USA | Feces | 101 | 302.81 | 394,906 |
| MSK464<sup>3</sup> | New York, USA | Feces | 101 | 287.72 | 394,569 |
| MSK465<sup>3</sup> | New York, USA | Feces | 101 | 252.57 | 397,063 |
| MSK466<sup>3</sup> | New York, USA | Feces | 101 | 304.11 | 395,274 |
| MSK606<sup>4</sup> | New York, USA | Blood | 101 | 344.82 | 352,155 |
| MSK607<sup>4</sup> | New York, USA | Colon | 101 | 358.94 | 352,121 |
| MSK798<sup>4</sup> | New York, USA | Blood | 101 | 394.31 | 351,303 |
| MSK801<sup>4</sup> | New York, USA | Blood | 101 | 413.37 | 351,401 |

<sup>1</sup> (Oh *et al.* 2019)
<sup>2</sup> (Pryszcz *et al.* 2015)
<sup>3</sup> (Zhai *et al.* 2020). MSK403-466 were isolated from a single patient.
<sup>4</sup> This study. MSK606, 607, 798 and 801 were isolated from two patients.

**Table S2.** Comparison of Illumina and minION assemblies of *C. metapsilosis* MSK414.

|  | **Illumina (SPAdes)** | **minION (Canu)** |
| --- | --- | --- |
| **Total number of contigs** | 13,527 | 45 |
| **Total length** | 26,170,059 | 27,138,054 |
| **Largest contig** | 115,451 | 3,141,946 |
| **N50** | 22,429 | 1,780,562 |
| **L50** | 337 | 6 |

**Table S3.** Assignment of contigs in *C. metapsilosis* MSK414 to haplotypes.

| **ATCC**<br>**96143**<br>**contig** | **MSK414 parent A** |  | **MSK414 parent C** |  |
| --- | --- | --- | --- | --- |
| 1.1<sup>1</sup><br>3,752,469 | tig3<sup>ct</sup> |  | tig1<sup>tct</sup> |  |
|  | Contig length | 2,603,961 | Contig length | 3,149,771 |
|  | Total length of all alignments | 2,367,064 | Total length of all alignments | 2,798,055 |
|  | Average identity | 94.81 | Average identity | 92.43 |
|  | **Identical bases** | **2,244,213** | **Identical bases** | **2,586,242** |
|  | tig11866<sup>t</sup> |  |  |  |
|  | Contig length | 488,734 |  |  |
|  | Total length of all alignments | 479,695 |  |  |
|  | Average identity | 95.75 |  |  |
|  | **Identical bases** | **459,308** |  |  |
|  | tig29<sup>tct</sup> |  | tig16<sup>tct</sup> |  |
|  | Contig length | 1,749,045 | Contig length | 1,784,933 |
|  | Total length of all alignments | 797,592 | Total length of all alignments | 775,958 |
|  | Average identity | 94.61 | Average identity | 91.86 |
|  | **Identical bases** | **754,602** | **Identical bases** | **712,795** |
| 2.1<br>2,105,923 | tig6<sup>tct</sup> |  | tig11867<sup>tct</sup> |  |
|  | Contig length | 2,096,600 | Contig length | 2,104,178 |
|  | Total length of all alignments | 2,010,934 | Total length of all alignments | 1,980,847 |
|  | Average identity | 94.64 | Average identity | 92.13 |
|  | **Identical bases** | **1,903,148** | **Identical bases** | **1,824,954** |
| 3.1<sup>1</sup><br>1,700,800 | tig10<sup>tct</sup> |  | tig11878<sup>tct</sup><br>MTLa |  |
|  | Contig length | 1,950,477 | Contig length | 1,289,153 |
|  | Total length of all alignments | 1,631,051 | Total length of all alignments | 729,610 |
|  | Average identity | 94.1 | Average identity | 91.70 |
|  | **Identical bases** | **1,534,819** | **Identical bases** | **669,052** |
|  |  |  | tig00011870 |  |
|  |  |  | Contig length | 954,831 |
|  |  |  | Total length of all alignments | 859,421 |
|  |  |  | Average identity | 93.94 |
|  |  |  | **Identical bases** | **807,340** |
| 4.1<br>1,367,707 | tig11876<sup>tct</sup> |  | tig39<sup>tct</sup> |  |
|  | Contig length | 1,338,872 | Contig length | 1,386,788 |
|  | Total length of all alignments | 1,287,149 | Total length of all alignments | 1,267,915 |
|  | Average identity | 94.59 | Average identity | 92.07 |
|  | **Identical bases** | **1,217,514** | **Identical bases** | **1,167,369** |
| 5.1<sup>1</sup><br>1,080,224 | tig11881<sup>tct</sup><br>MTLalpha<br>rDNA |  | tig11874<sup>t</sup> |  |
|  | Contig length | 1,095,871 | Contig length | 635,722 |
|  | Total length of all alignments | 1,013,327 | Total length of all alignments | 497,399 |
|  | Average identity | 95.41 | Average identity | 92.30 |
|  | **Identical bases** | **966,815** | **Identical bases** | **459,099** |
|  |  |  | tig00011878<sup>tct</sup><br>MTLa |  |
|  |  |  | Contig length | 1,289,153 |
|  |  |  | Total length of all alignments | 498,349 |
|  |  |  | Average identity | 91.60 |
|  |  |  | **Identical bases** | **456,488** |
| 6.1<sup>1</sup><br>903,567 | tig29<sup>tct</sup> |  | tig16<sup>tct</sup> |  |
|  | Contig length | 1,749,045 | Contig length | 1,784,933 |
|  | Total length of all alignments | 634,361 | Total length of all alignments | 634,288 |
|  | Average identity | 94.7 | Average identity | 91.71 |
|  | **Identical bases** | **600,740** | **Identical bases** | **581,706** |
|  | tig3<sup>ct</sup> |  | tig1<sup>tct</sup> |  |
|  | Contig length | 2,603,961 | Contig length | 3,149,771 |
|  | Total length of all alignments | 216,182 | Total length of all alignments | 213,770 |
|  | Average identity | 94.46 | Average identity | 92.45 |
|  | **Identical bases** | **204,206** | **Identical bases** | **197,630** |
| 7.1<br>704,064 | tig11880<sup>tct</sup> |  | tig11884<sup>tct</sup><br>rDNA |  |
|  | Contig length | 1,017,382 | Contig length | 904,741 |
|  | Total length of all alignments | 655,192 | Total length of all alignments | 653,023 |
|  | Average identity | 93.79 | Average identity | 92.93 |
|  | **Identical bases** | **614,505** | **Identical bases** | **606,854** |
| 8.1<br>677,266 | tig64<sup>tct</sup> |  | tig60<sup>tct</sup> |  |
|  | Contig length | 798,529 | Contig length | 802,845 |
|  | Total length of all alignments | 592,388 | Total length of all alignments | 564,780 |
|  | Average identity | 94.52 | Average identity | 91.63 |
|  | **Identical bases** | **559,925** | **Identical bases** | **517,508** |
| All contigs | **Total alignment length across all contigs** | 11,684,935 | **Total alignment length across all contigs** | 11,473,415 |
|  | **Identical bases across all contigs** | 11,059,795 | **Identical bases across all contigs** | 10,587,038 |
|  | **Average identity across all contigs** | **94.65%** | **Average identity across all contigs** | **92.27%** |

<sup>t</sup> Contains a telomere at one or both ends.
<sup>c</sup> Contains a centromere.
<sup>1</sup>Where contigs resulted from a translocation between parts of two different contigs in the ATCC 96143 assembly, the sequence of the relevant contig was aligned to both contigs in the ATCC 96143 assembly. The average nucleotide identity for each alignment is shown separately. The average identity across all contigs was calculated as the total number of identical bases in all alignments divided by the total length of all alignments across all contigs. There appears to be an assembly error at the tip of scaffold 6 in the assembly of *C. metapsilosis* ATCC 96143 (Oh *et al.* 2019) (see Figure S2).

